# Proteasome-dependent degradation and nucleus-vacuole junctions sustain proteostasis during acute glucose starvation

**DOI:** 10.64898/2026.04.22.720209

**Authors:** Mihaela Pravica, Dina Franić, Matko Bazdan, Dominik Guzalić, Antonio Bedalov, Mirta Boban

**Affiliations:** University of Zagreb School of Medicine, Croatian Institute for Brain Research, Zagreb, Croatia; Fred Hutchinson Cancer Center, Seattle, USA

**Keywords:** ATP, glucose, energy metabolism, proteasome, ubiquitin, vacuole, yeast, protein quality control, nucleus-vacuole junction, membrane-contact sites

## Abstract

How protein quality control is maintained during acute metabolic stress remains poorly understood. In budding yeast, abrupt glucose depletion rapidly lowers ATP levels and leads to the formation of chaperone-containing inclusions, suggesting that ATP-dependent degradation of misfolded proteins may be compromised when energy becomes limiting. Here we find that selective degradation of misfolded proteins remains active during acute glucose starvation despite reduced cellular ATP levels. Using model misfolded substrates in yeast *Saccharomyces cerevisiae*, we show that misfolded proteins continue to be efficiently degraded throughout both early and late phases of acute glucose depletion. This degradation requires the proteasome and depends on its functional 19S regulatory particle, indicating that ATP-dependent proteasomal activity persists during metabolic stress. We further find that nucleus–vacuole junctions (NVJs) promote efficient degradation during prolonged glucose starvation, revealing a role for organelle contact sites in supporting proteostasis under energy limitation. Together, these findings indicate that cells preserve proteasome-mediated proteostasis during acute glucose starvation, while NVJ membrane contact sites help sustain degradation capacity when metabolic resources are scarce.

## INTRODUCTION

To maintain proteostasis, eukaryotic cells rely on a complex network of molecular pathways collectively referred to as the protein quality control (PQC) system. This system prevents the accumulation of misfolded proteins through protein folding, selective degradation, and sequestration into specialized compartments (1–5). Many of these processes are ATP-dependent, including protein folding, ubiquitination, and selective degradation (6), suggesting that PQC is vulnerable to fluctuations in cellular energy levels. Yet, the mechanisms by which cells preserve proteostasis during metabolic stress and energy limitation remain poorly understood.

Cells often face fluctuating environments in which nutrient availability changes over time. The budding yeast *Saccharomyces cerevisiae* uses glucose as its preferred carbon source and, when glucose is available, generates ATP primarily through glycolysis coupled to fermentation, even in the presence of oxygen (7). The cellular response to glucose scarcity depends on the rate at which glucose is depleted from the medium. Gradual glucose exhaustion during yeast growth is associated with regulated entry into quiescence and a metabolic shift towards respiration and the use of ethanol, which is produced earlier through glycolysis-coupled fermentation (8–10). While yeast cells eventually adapt to acute glucose starvation by switching to alternative energy-generating pathways such as mitochondrial respiration, β-oxidation and macroautophagy (11–15), ATP levels stabilize at only around 60 % of pre-starvation levels (13, 16, 17), indicating that cells remain in a reduced energy state. How cells maintain proteostasis during acute glucose starvation, a condition associated with severe energy stress, remains unclear.

The main pathway for degradation of misfolded proteins in the cell is regulated, ATP-dependent proteolysis mediated by the ubiquitin proteasome system (UPS) (5, 18, 19). Protein degradation by the UPS requires ATP at multiple steps, including ubiquitin activation by the E1 enzyme (20) and activity of proteasomal ATPases that mediate gate opening (21) and substrate unfolding (22). Recent studies have shown that acute glucose depletion leads to the formation of cytosolic inclusions containing Hsp70 chaperones and Hsp104, an ATP-dependent disaggregase widely used as a marker of misfolded protein aggregates (23, 24, 25). The appearance of Hsp104-positive inclusions suggested that ATP-dependent degradation of misfolded proteins might be compromised when energy becomes limiting.

Glucose withdrawal also results in expansion of nucleus-vacuole junctions (NVJs), a membrane contact site formed by the interaction of Nvj1 and Vac8 (26–29). The NVJ is the site of piecemeal microautophagy of the nucleus (PMN) (27, 30), and accumulating evidence suggests that it contributes to PQC of misfolded proteins through the delivery of protein inclusions to the vacuole (31–33).

In this study, we investigated selective degradation of misfolded proteins in yeast cells subjected to acute glucose depletion. We show that glucose-starved cells maintain selective degradation of misfolded proteins and that this process requires functional 26S proteasomes. Furthermore, we identify the NVJ as a key contributor to sustained degradation during prolonged starvation, acting independently of core Atg1-dependent autophagy. Together, our findings reveal that cells preserve proteostasis under acute energy limitation and uncover a role for organelle contact sites in supporting degradation capacity during metabolic stress.

## RESULTS

### Cells subjected to an abrupt glucose depletion sustain selective degradation of misfolded proteins despite reduced ATP levels

To investigate selective degradation of misfolded proteins in yeast cells subjected to acute glucose depletion, exponentially growing yeast cells were pre-cultured in glucose-replete medium containing 2% glucose and then transferred to medium containing 0.2% or 0.02% glucose, followed by incubation for the indicated periods. Control cells were transferred to medium containing 2% glucose and incubated for the same duration.

As a first step, we measured intracellular ATP levels in cells subjected to acute glucose depletion (**Figure 1A**). In cells incubated in medium containing 0.02% glucose, ATP levels declined sharply within 15 min of glucose depletion and were partially recovered by 90 min, reaching approximately 60% of control levels. ATP levels measured after 4 h remained at a similar level. By contrast, in cells incubated in 0.2% glucose, ATP levels were fully recovered by 90 min to values comparable to those of control cells maintained in 2% glucose. As an additional reference condition, cells were incubated with iodoacetic acid (IAA), which causes severe ATP depletion (34). Together, these data are consistent with earlier studies showing that acute glucose depletion results in an initial sharp decline in ATP levels, followed by a partial recovery to around 60% of control levels (13, 16, 17).

**Figure 1.**
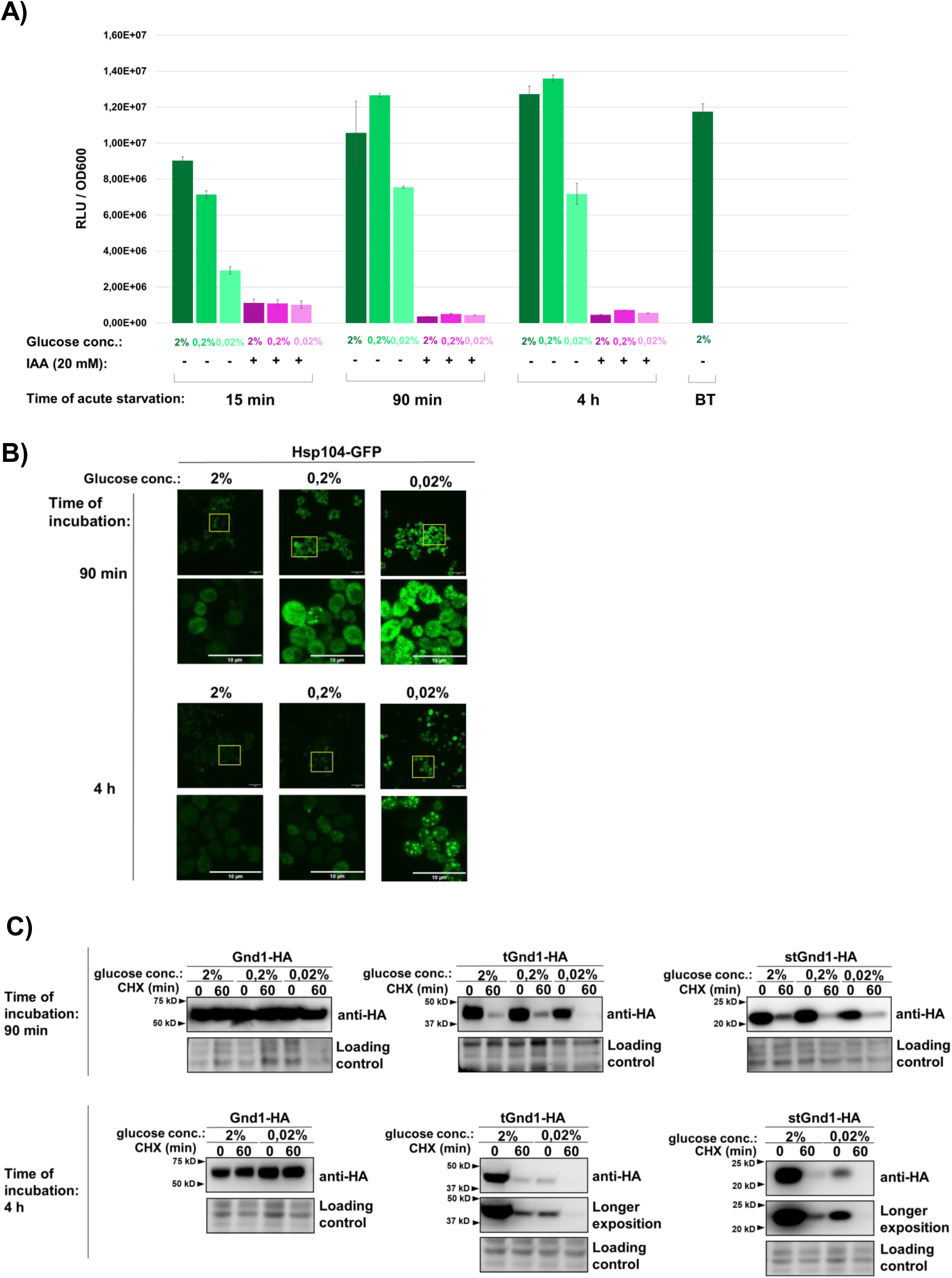
Acute glucose starvation triggers ATP decline and spatial reorganization of the disaggregase Hsp104 while maintaining selective degradation of model misfolded proteins tGnd1-HA and stGnd1-HA. **A) Cellular ATP levels during acute glucose starvation.** Logarithmically growing cells of the wild type yeast strain (BY4741) were shifted from media containing 2% glucose to media containing 2%, 0.2%, or 0.02% glucose and incubated for the indicated time period. Relative ATP levels were measured using a bioluminescence-based BacTiter assay, and bioluminescence (relative luminescence units, RLU) was normalized to optical density of cells (RLU/OD_600_). Data show measurements taken at 15 min, 90 min, and 4 h of acute glucose depletion. Parallel cultures were supplemented with iodoacetic acid (IAA, 20 mM) to verify the metabolic specificity of the bioluminescent signal. “BT” indicates measurements taken immediately prior to the acute glucose depletion (“before treatment”). Data are presented as average values ± SD RLU/ OD_600_ (n = 2). **B) Hsp104-GFP accumulates in foci during acute glucose starvation.** Yeast cells expressing Hsp104-GFP were exponentially grown in the medium containing 2% glucose, and either maintained under glucose replete conditions (2% glucose) or subjected to acute glucose depletion (0.2% or 0.02% glucose) for the indicated time periods (90 min, 4 h), prior to imaging by confocal microscopy. Shown is the whole z-stack. Scale bar, 10 µm. Representative images are shown. **C) Model misfolded proteins tGnd1-HA and stGnd1-HA are selectively degraded during early and late phases of acute glucose starvation.** Representative immunoblots showing the protein stability of native Gnd1-HA and model misfolded proteins tGnd1-HA and stGnd1-HA. Yeast cells expressing the indicated constructs were grown as in (A) and shifted to media containing either 2%, 0.2%, or 0.02% glucose for 90 min (early phase) or 4 h (late phase). Protein stability was assessed by cycloheximide (CHX) chase at the indicated time points, followed by Western blot (anti-HA). Stain-free total protein is shown as a loading control.

Previous studies have shown that acute glucose depletion promotes the formation of Hsp104-GFP inclusions (22, 24). Hsp104 is an ATP-dependent disaggregase commonly used as a reporter of protein aggregation (23). To determine whether our growth conditions induce a similar response, we monitored the subcellular localization of Hsp104-GFP in cells subjected to acute glucose depletion (**Figure 1B**). Consistent with previous findings, Hsp104-GFP displayed a diffuse cytosolic localization under glucose-replete conditions (2% glucose; **Figure 1B**). In contrast, 90 min of glucose starvation (0.2% or 0.02% glucose) induced the formation of Hsp104-GFP foci (**Figure 1B**, upper panel), whereas larger and more prominent inclusions were observed after 4 h of starvation (**Figure 1B**, lower panel). Taken together, these results show that glucose depletion induces the formation of Hsp104-GFP foci, consistent with previous studies (24, 25).

The formation of Hsp104-GFP inclusions during glucose starvation suggested sequestration of misfolded proteins and raised the possibility that their clearance might be compromised. To test this, we examined two well-characterized model misfolded proteins, tGnd1 (truncated Gnd1) and stGnd1 (small truncated Gnd1), which are C-terminally truncated variants of the yeast 6-phosphogluconate dehydrogenase Gnd1 generated by premature stop codons in the *GND1* gene (35). We asked whether acute glucose depletion affects degradation of tGnd1 and stGnd1 by assessing their stability in cycloheximide chase assays and comparing it with that of the native protein Gnd1. Whereas native Gnd1-HA remained stable, the misfolded proteins tGnd1-HA and stGnd1-HA were degraded in cells incubated in medium containing 2%, 0.2%, or 0.02% glucose during both the early (90 min) and prolonged (4 h) phases of starvation (**Figure 1C**). These results indicate that cells subjected to acute glucose depletion maintain selective degradation of misfolded proteins.

### Sequestration of misfolded proteins under acute glucose depletion remains substrate-specific

Next, we visualized misfolded proteins during acute glucose starvation using mNeonGreen-tagged variants of tGnd1 and stGnd1 (**Figure 2**). We first verified that the mNeonGreen-tagged variants were degraded at rates comparable to those of the untagged proteins. Biochemical analysis confirmed degradation of both mNeonGreen-tagged tGnd1 and stGnd1 under acute glucose starvation conditions (**Figure 2A**, **C**). In glucose-replete medium (2% glucose), mNeonGreen-tGnd1-HA showed diffuse cytosolic localization, with approximately 25% of cells containing one or two inclusions (**Figure 2B**). A similar localization pattern was observed under moderate glucose depletion (0.2% glucose) (**Figure 2B**), whereas severe glucose depletion (0.02% glucose) resulted in a weaker diffuse signal, a modest increase in the overall fraction of cells containing inclusions (to approximately 33%) and a reduced percentage of cells with two or more inclusions (**Figure 2B**). The weaker diffuse signal and lower fraction of cells with two or more inclusions likely reflect reduced expression of mNeonGreen-tGnd1-HA under this condition, consistent with the Western blot analysis (**Figure 2A**). In agreement with continued protein degradation detected by immunoblotting, cells harvested 60 min after cycloheximide addition showed a clear reduction in fluorescence signal by microscopy (**Figure 2B**). Taken together, these results indicate that severe glucose reduction leads to only a modest increase in mNeonGreen-tGnd1 inclusion formation, whereas its degradation remains largely unchanged across conditions.

**Figure 2.**
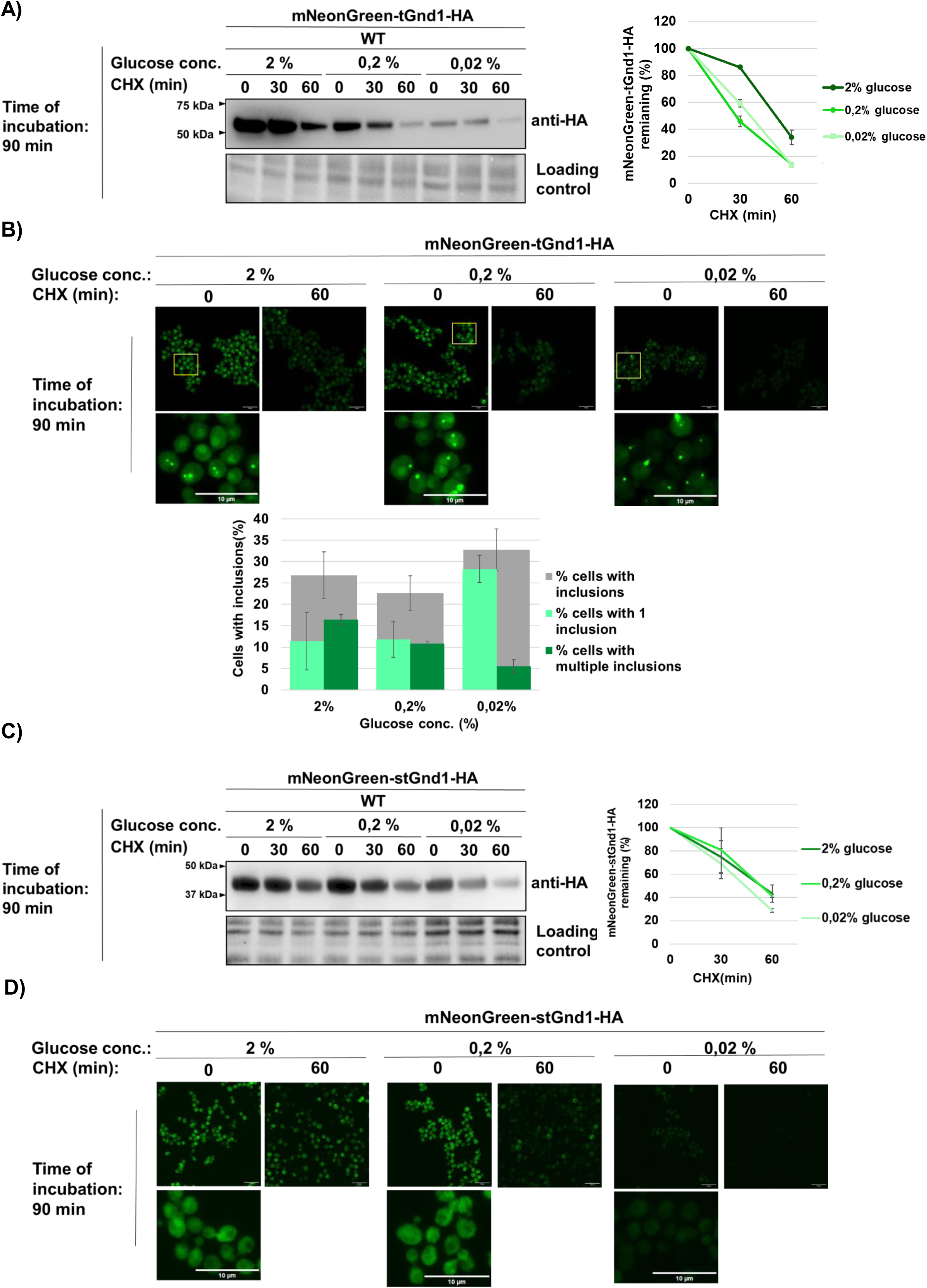
Acute glucose starvation preserves substrate-specific sequestration of misfolded proteins into inclusions. Wild type cells expressing indicated model misfolded proteins were grown under glucose-replete conditions (medium containing 2% glucose) to the logarithmic growth phase and then subjected to 90 min of acute glucose depletion (0.02% glucose) or glucose-replete conditions as a control (2% glucose). **(A, C)** Representative immunoblots showing protein stability of mNeonGreen-tGnd1-HA (A) or mNeonGreen-stGnd1-HA (C). Protein stability was monitored by cycloheximide chase at the indicated time points, followed by Western blot (anti-HA). Stain-free total protein is shown as a loading control. Graphs represent mNeonGreen-tGnd1-HA or mNeonGreen-stGnd1-HA protein levels as a percentage of the protein present at the time point 0 min after the addition of cycloheximide. Data are presented as average ± SD from 2 independent experiments. **(B, D)** Representative confocal fluorescence microscopy images of yeast cells expressing mNeonGreen-tGnd1-HA (B) or mNeonGreen-stGnd1-HA, cultured as above. Images represent the whole z-stack. Scale bar, 10 µm. The graph in (B) shows the percentage of cells with inclusions of mNeonGreen-tGnd1-HA (grey), and the percentage of cells harboring one (light green) or multiple (dark green) inclusions. Data are presented as average ± SD from 2 independent experiments. At least 140 cells were counted per cell culture. mNeonGreen-stGnd1-HA retains a diffuse localization (D).

We also examined intracellular distribution of mNeonGreen-tagged stGnd1 (**Figure 2D**), a shorter variant of Gnd1. This protein exhibited a largely uniform intracellular distribution both under glucose-replete conditions (2% glucose) (**Figure 2D**) and after 90 min of acute glucose depletion (0.2% and 0.02% glucose) (**Figure 2D**). The data indicate that inclusion formation is not a general strategy for misfolded protein management during acute glucose starvation.

To determine whether selective degradation represents a general cellular strategy for managing misfolded proteins during acute glucose depletion, rather than an effect specific to the Gnd1 variants, we examined the degradation of an additional model misfolded protein, NBD2*-HA (**Supplementary Figure S1**), a well-characterized truncated variant of the yeast ABC transporter Ste6 (36). Consistent with the previous study, under glucose-replete conditions (2% glucose), the misfolded protein NBD2*-HA was degraded, whereas the corresponding native protein, NBD2-HA, remained stable. A similar pattern was observed in cells incubated for 4 h in medium containing 0.02% glucose (**Supplementary Figure S1**), indicating that selective degradation of misfolded proteins is maintained during acute glucose starvation. In addition to a weak diffuse signal throughout the cell, mNeonGreen-tagged NBD2* formed visible inclusions in around 15% of cells under all tested conditions (2%, 0.2%, and 0.02% glucose) (**Supplementary Figure S2A**). Biochemical analysis using cycloheximide chase and immunoblotting showed that HA-mNeonGreen-NBD2* was degraded similarly under all tested conditions, including acute glucose starvation (**Supplementary Figure S2B**). Together, these data show that acute glucose depletion does not lead to a substantial increase in sequestration of misfolded proteins tGnd1 and NBD2* into inclusions. Instead, cells retain the capacity to selectively degrade these proteins. Moreover, sequestration of misfolded proteins under acute glucose depletion remains substrate-specific rather than serving as a general strategy for misfolded protein management.

### Selective degradation of misfolded proteins during acute glucose starvation requires proteasomal activity

Having established that misfolded proteins are efficiently cleared during acute glucose starvation, we next asked whether this process depends on the proteasome by assessing the stability of tGnd1-HA and stGnd1-HA in cells treated with the proteasome inhibitor bortezomib. These experiments were performed in the *pdr5Δ* mutant, which lacks the Pdr5 efflux pump and is therefore sensitive to the drug (37). Whereas both misfolded proteins were rapidly degraded in control cells during the early (90 min) and late (4 h) phases of acute glucose depletion (0.02% glucose), bortezomib treatment markedly stabilized both proteins under all tested conditions, indicating that their degradation is proteasome-dependent (**Figure 3A**). We further examined a third model misfolded protein, NBD2*-HA, which was similarly stabilized by bortezomib after 4 h of acute glucose starvation (**Supplementary Figure S3**). In contrast, the native protein NBD2-HA remained stable regardless of proteasome inhibition. Together, these results establish that selective degradation of misfolded proteins during acute glucose starvation requires proteasomal activity.

**Figure 3.**
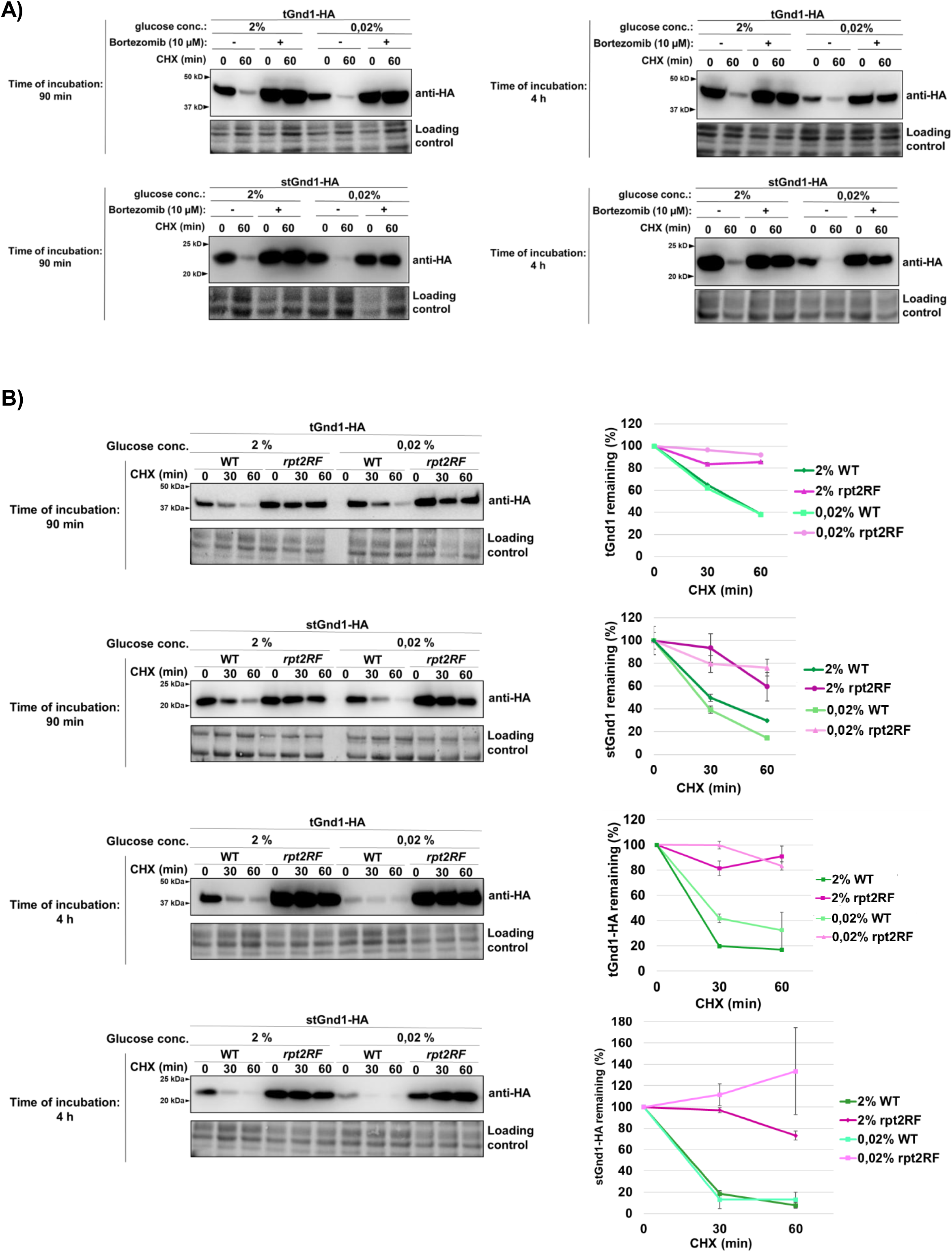
Degradation of tGnd1-HA and stGnd1-HA during acute glucose depletion depends on the 26S proteasome. Cells expressing indicated model misfolded proteins were grown in the medium containing 2% glucose to the logarithmic growth phase and then subjected to 90 min or 4 h of acute glucose depletion (0.02% glucose) or glucose-replete conditions as a control (medium containing 2% glucose). Protein stability was assessed by cycloheximide chase (CHX, 100 µg/mL); cells were harvested at the indicated time points followed by Western blot analysis (anti-HA). Stain-free total protein is shown as a loading control. **A) Degradation of tGnd1-HA and stGnd1-HA during both the early and late phases of acute glucose starvation depends on proteasomal proteolytic activity.** Representative immunoblots showing protein stability of the indicated model misfolded proteins in the presence or absence of the proteasome inhibitor bortezomib. Drug-sensitive *pdr5Δ* mutant strains expressing indicated model misfolded proteins were used. Bortezomib was added at the final concentration of 10 µM 30 min prior to the addition of cycloheximide. **B) Degradation of tGnd1-HA and stGnd1-HA requires a functional Rpt2 ATPase of the 19S regulatory proteasome particle.** Representative immunoblots showing protein stability of tGnd1-HA and stGnd1-HA expressed in *rpt2RF* mutant (YP55) and the respective wild type (WT) strain (MPY100) from centromeric plasmids (pMB214 and pMB215), grown at the temperatures indicated in the section Experimental procedures. Graphs show tGnd1-HA and stGnd1-HA protein levels as a percentage of the protein present at the time point 0 min after the addition of cycloheximide. Average values and standard deviation are shown (n = 2).

Given that acute glucose starvation leads to a decline in cellular ATP levels (13, **Figure 1A**), we asked whether the 19S regulatory particle, which mediates ubiquitin recognition and ATP-dependent substrate unfolding (38), remains required for misfolded protein degradation under these conditions. Using the *rpt2RF* mutant strain, which expresses a defective ATPase Rpt2, we found that both tGnd1-HA and stGnd1-HA were significantly stabilized during the early (90 min) and late (4 h) phases of starvation (0.02% glucose), whereas they were efficiently cleared in wild-type cells (**Figure 3B**). This result indicates that the proteasome 19S regulatory particle remains required for degradation of misfolded proteins during acute glucose starvation, despite the marked reduction in ATP availability (**Figure 1A**).

### Degradation of misfolded proteins becomes increasingly dependent on the nucleus-vacuole junction (NVJ) during the later phases of acute glucose starvation

Our recent work showed that efficient degradation of specific misfolded proteins in quiescence, a cellular state induced by gradual glucose exhaustion, requires functional nucleus-vacuole junctions (NVJs) (33). NVJs are membrane contact sites formed through direct interaction between Nvj1 and Vac8, integral membrane proteins of the outer nuclear membrane and the vacuole, respectively. Since NVJs undergo substantial expansion and proteomic remodeling upon glucose withdrawal (27, 33), we asked whether they also contribute to the clearance of misfolded proteins during acute glucose depletion.

To address this, we analyzed the stability of tGnd1-HA in a *nvj1Δ* strain, which is defective in NVJ formation (26). Under glucose-replete conditions (2% glucose), degradation of tGnd1-HA was independent of functional NVJ, consistent with our previous findings (33). At an early time point after glucose depletion (90 min in 0.02% glucose), degradation of tGnd1-HA in *nvj1Δ* cells was comparable to that in wild-type cells (**Figure 4**, left, upper panel), indicating that tGnd1 degradation rate remains unimpaired. In contrast, after prolonged glucose starvation (4 h), tGnd1-HA degradation was slower in *nvj1Δ* cells (**Figure 4**, left, lower panel), indicating that functional NVJs are required for efficient tGnd1 clearance during later stages of acute glucose starvation. By contrast, degradation of another model misfolded protein, stGnd1-HA, was unaffected by loss of Nvj1 at either time point (**Figure 4**, right, upper and lower panel), suggesting that the contribution of NVJs to misfolded protein clearance is substrate-specific. Together, these findings indicate that sustained clearance of specific substrates during prolonged metabolic stress requires functional NVJs.

**Figure 4.**
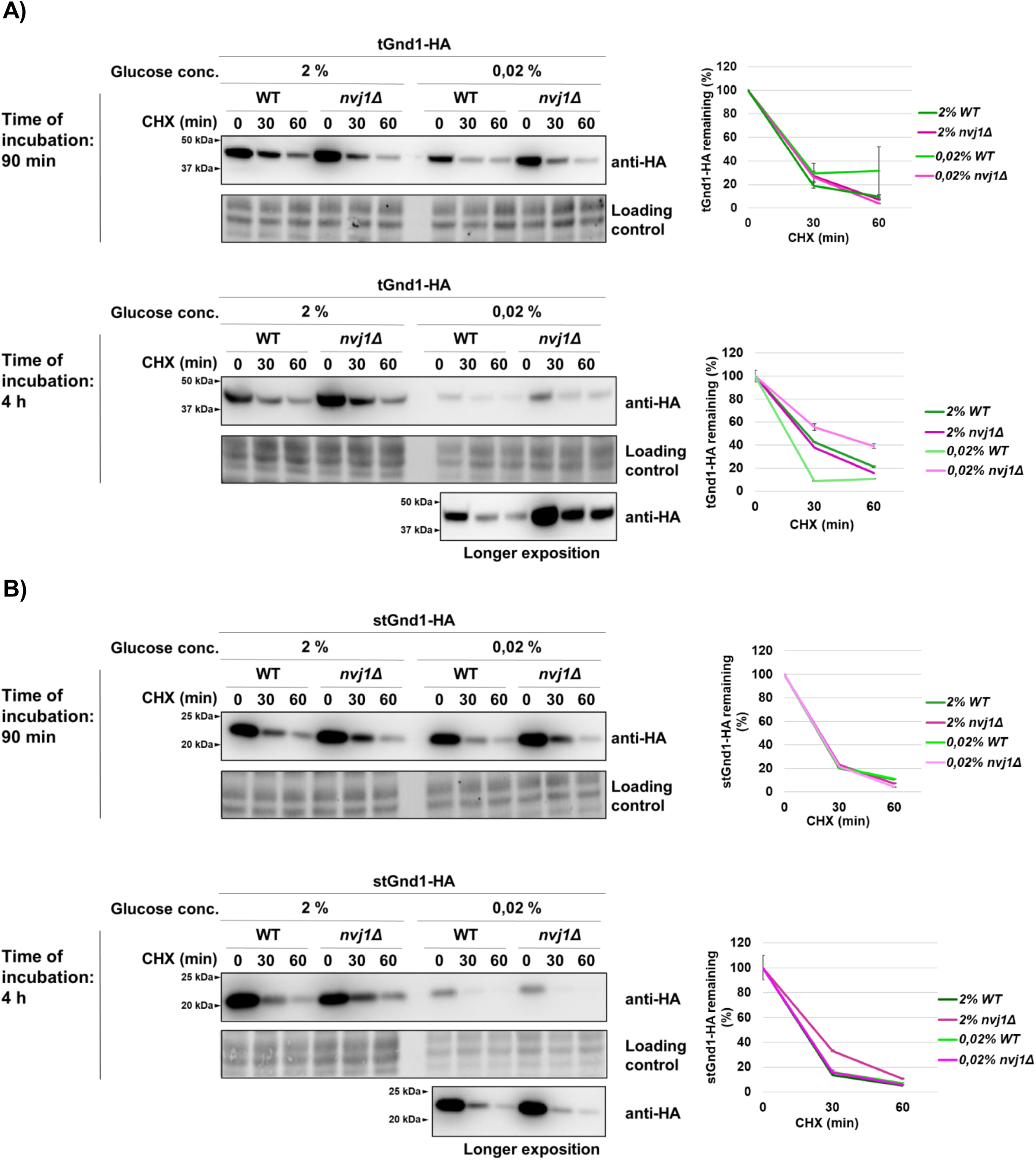
Misfolded protein tGnd1-HA, but not stGnd1-HA, is stabilized in the nucleus-vacuole junction mutant *nvj1Δ* during prolonged acute glucose starvation. Representative immunoblots of tGnd1-HA and stGnd1-HA. Logarithmically growing cells of wild-type (WT) and *nvj1Δ* strains expressing tGnd1-HA or stGnd1-HA were subjected to acute glucose starvation (0.02% glucose) for 90 min (early phase) or 4 h (late phase), or maintained in glucose-replete medium (2% glucose). Protein stability was monitored by cycloheximide chase assay (CHX, 100 µg/mL) and protein levels were analyzed by Western blot (anti-HA). Total proteins were visualized using stain-free technology (BioRad). Graphs represent tGnd1-HA and stGnd1-HA protein levels as a percentage of the protein present at the time point 0 min after the addition of cycloheximide. Average values and standard deviation are shown (n=2).

Next, we asked whether the role of NVJ becomes apparent already during the early phase of acute glucose starvation (90 min) when proteasome activity is compromised (**Figure 5**). Compared with *NVJ1*-wild type cells treated with bortezomib, bortezomib-treated *nvj1Δ* mutant cells exhibited further stabilization of tGnd1-HA in glucose depletion (**Figure 5**, lower panel). This additive effect of proteasome inhibition and NVJ loss indicates that the proteasome and NVJ represent two distinct pathways that function in parallel. Furthermore, these results suggest that even during the early phase of acute glucose starvation (90 min in 0.02% glucose), cells rely on NVJs for clearance of misfolded proteins, although to a lesser extent than during later stages of starvation.

**Figure 5.**
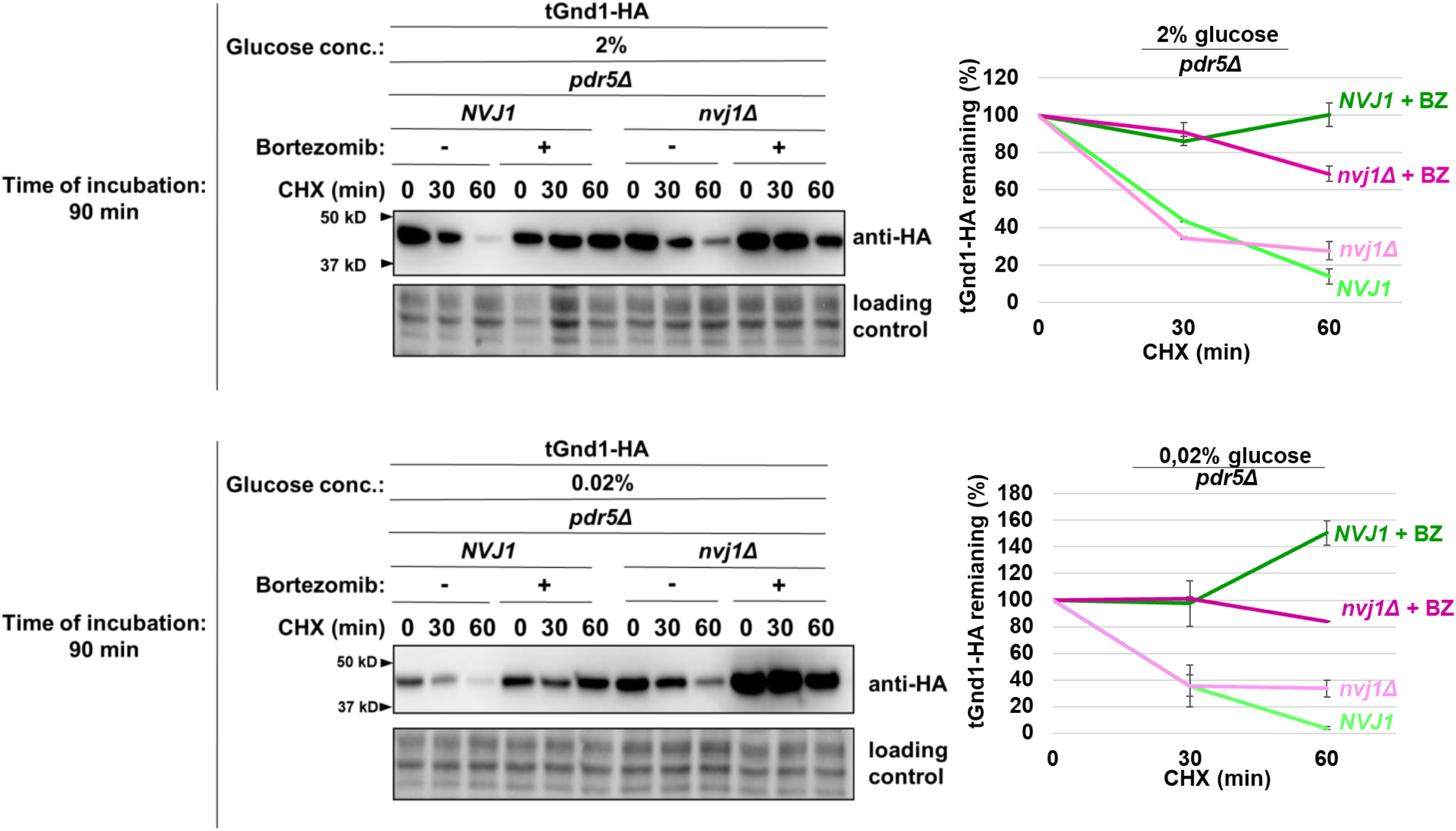
Additional stabilization of tGnd1-HA upon proteasome inhibition in *nvj1Δ* cells indicates that nucleus-vacuole junctions and the proteasome define parallel clearance pathways during acute glucose depletion. Representative immunoblots of tGnd1-HA expressed in drug-sensitive *pdr5Δ* strains, either *NVJ1* wild type (*NVJ1 pdr5Δ*) or *nvj1Δ* deletion mutant (*nvj1Δ pdr5Δ*), subjected to acute glucose depletion. Logarithmically growing cells were shifted from 2% glucose-containing medium to glucose starvation conditions (0.02% glucose) for a time period of 90 min, followed by proteasome inhibition with 10 μM bortezomib for 30 min prior to cycloheximide (CHX, 100 μg/ml) addition. Protein levels at the indicated time points were assessed by Western blot (anti-HA). Stain-free total protein is shown as a loading control. Graphs represent tGnd1-HA protein levels as a percentage of the protein present at the time point 0 min. Average values and standard deviation are shown (n=2).

### Clearance of misfolded proteins during acute glucose starvation does not require Atg1-dependent autophagy

Reports on the glucose starvation-induced autophagy are controversial (11, 12, 14, 15, 39, 40). To assess bulk autophagy induction under our experimental conditions, we used the GFP-Atg8 processing assay (41). After 4 h of acute glucose deprivation (0.02% glucose), the free GFP signal was very weak and similar to that observed under glucose-replete conditions (2% glucose) (**Figure 6A**), indicating little or no autophagy induction. Moreover, GFP-Atg8 processing was similar in wild-type and *atg1Δ* cells, further supporting the absence of substantial Atg1-dependent autophagic flux. These findings are consistent with previous reports showing that acute glucose starvation does not induce canonical autophagy (11, 12, 14, 15).

**Figure 6.**
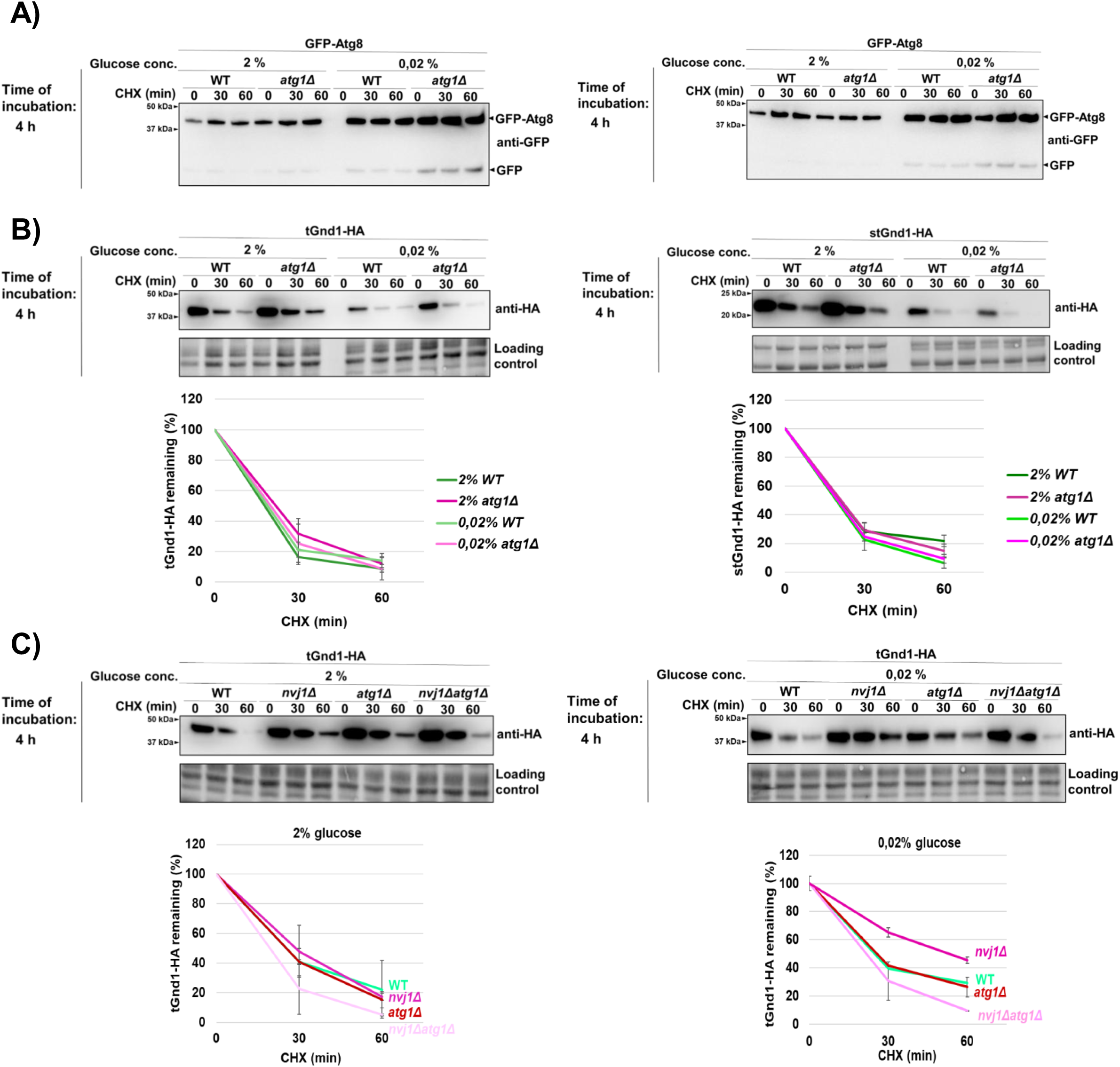
Degradation of tGnd1-HA and stGnd1-HA during acute glucose depletion is independent of Atg1. Logarithmically growing cells were subjected to acute starvation (0.02% glucose) for 4 h, or maintained in glucose-replete medium (2% glucose). Protein degradation was monitored by cycloheximide chase (CHX, 100 µg/mL), by harvesting cells at indicated time points after the addition of cycloheximide and the analysis of protein levels by Western blot. Stain-free total protein is shown as a loading control. **A)** GFP-Atg8 processing assay assessing autophagy induction in wild type (WT) and *atg1Δ* mutant under acute glucose starvation conditions. Total cell lysates as shown below in (B) were analyzed by Western blot using anti-GFP antibodies. The migration of free GFP species and GFP-Atg8 fusion protein is indicated. **B)** Representative immunoblots (anti-HA) showing the stability of tGnd1-HA and stGnd1-HA in WT and autophagy-deficient strain *atg1Δ*. Graphs represent indicated protein levels as a percentage of the protein present at the time point 0 min after the addition of cycloheximide. Average values and standard deviation are shown (n=2). **C)** Representative immunoblots (anti-HA) of tGnd1-HA expressed in wild-type (WT), *nvj1Δ* and *atg1Δ* single mutants, and double *nvj1Δ atg1Δ* mutant strain. Graphs represent tGnd1-HA protein levels as a percentage of the protein present at the time point 0 min. Average values and standard deviation are shown (n=2).

To test whether clearance of misfolded proteins during acute glucose starvation requires Atg1-dependent autophagy, we examined the stability of tGnd1-HA and stGnd1-HA in the *atg1Δ* mutant. Degradation of both tGnd1-HA and stGnd1-HA in cells subjected to acute glucose depletion (0.02% glucose for 4 hours) was unaffected in *atg1Δ* cells (**Figure 6B**), indicating that Atg1-dependent autophagy is not required for clearance of misfolded proteins under these conditions.

To determine whether a role for Atg1-dependent autophagy during acute glucose starvation becomes apparent when NVJ function is disrupted, we compared tGnd1-HA stability in the single mutants *nvj1Δ* and *atg1Δ* and in the double mutant *nvj1Δ atg1Δ* (**Figure 6C**). At the 4-h time point, when NVJ-dependent effects on tGnd1-HA clearance become evident (**Figure 4, lower panels**), the degradation of tGnd1-HA in the *nvj1Δ atg1Δ* double mutant was similar to that in the single mutants, with no additive stabilization observed (**Figure 6C**). Together, these findings indicate that Atg1-dependent autophagy does not contribute to clearance of misfolded proteins during acute glucose starvation.

## DISCUSSION

Acute glucose starvation poses a major metabolic stress, yet how cells preserve proteostasis under such conditions has remained unclear. Here, we show that the yeast *Saccharomyces cerevisiae* maintains selective degradation of misfolded proteins through the 26S proteasome during acute glucose depletion, demonstrating that this ATP-dependent PQC pathway remains functional despite severe nutrient stress. Although efficient clearance of misfolded proteins is sustained during both the early and prolonged phases of acute glucose starvation, nucleus-vacuole junctions (NVJs) make an increasingly important contribution during prolonged starvation.

Abrupt glucose withdrawal leads to an initial sharp drop in intracellular ATP levels, followed by partial recovery and stabilization at around 60% of pre-starvation levels (13, 16, 17) (**Figure 1 A**). Previous studies showed that acute glucose depletion leads to the formation of Hsp70– and Hsp104-positive inclusions, suggesting that ATP-dependent clearance of misfolded proteins may be transiently compromised when energy becomes limiting (24, 25). However, protein degradation during acute glucose starvation has not been directly investigated. In agreement with these reports, we also observed the formation of Hsp104-GFP foci during glucose depletion (**Figure 1 B**). However, despite this phenotype and reduced ATP levels, the model misfolded proteins examined in our study remained efficiently degraded, with only a modest increase in foci formation (**Figure 2**).

Our findings showing sustained 26S proteasome-dependent degradation of misfolded proteins indicate that acute glucose starvation does not abolish ATP-dependent degradation of PQC substrates. The data suggest that these ATP levels are sufficient to support continued degradation-mediated PQC, and that degradation-mediated PQC is a prioritized pathway for proteostasis maintenance. In line with this, the model misfolded protein stGnd1, which does not form foci in glucose replete conditions, also did not form foci during acute glucose depletion, suggesting that sequestration is not a general PQC strategy for managing misfolded proteins under this form of nutrient stress. More pronounced inclusion formation during acute glucose starvation, as observed for the misfolded protein Ubc9ts in a previous study, may reflect stronger and more sustained ATP depletion caused by incubation with 2-deoxy-D-glucose, a potent inhibitor of glycolysis (16, 17, 25, 42), or may instead be more characteristic of highly aggregation-prone proteins such as Rnq1 (24).

Our data further show that the contribution of NVJs becomes more prominent during prolonged acute starvation. At the earlier time point, NVJ contribution became apparent only upon proteasome inhibition by bortezomib (**Figure 5**). This observation indirectly suggests that the capacity of proteasome-mediated degradation becomes more limited at later stages of acute glucose starvation, thereby increasing reliance on NVJ. Furthermore, degradation of stGnd1, which does not form foci, was unaffected in the *nvj1Δ* mutant and remained fully proteasome-dependent. Thus, the NVJ requirement is clearly substrate-specific. This finding suggests that the increasing requirement for the NVJ in tGnd1 degradation during prolonged glucose starvation is unlikely to reflect a general decline in proteasome activity *per se*. Rather, another component of the UPS degradation pathway may become limiting. Identifying such factors will be an interesting open question for future studies. Possible candidates include ATP-dependent chaperones, whose reduced activity during prolonged glucose depletion could result in a decreased solubility or proteasome accessibility of specific misfolded proteins such as tGnd1, thereby creating an additional requirement for NVJ function. Further open questions concern the precise mechanism by which NVJs mediate misfolded protein clearance, and their possible interplay with proteasomal degradation.

Our results do not support a major role for Atg1-dependent autophagy in the degradation of misfolded proteins during acute glucose starvation. Degradation of misfolded substrates was unaffected in *atg1Δ* cells, GFP-Atg8 processing indicated little or no induction of bulk autophagy, and combined loss of Atg1 and Nvj1 did not further impair tGnd1 clearance beyond the *nvj1Δ* phenotype alone (**Figure 6**). Thus, the NVJ contribution to misfolded protein clearance during acute glucose starvation appears to be independent of the canonical Atg1-dependent autophagy, although we cannot exclude the involvement of other forms of autophagy. In line with our findings, a previous study showed that intracellular ATP levels in the *atg2Δ* mutant were unaffected during the first hour of acute glucose starvation, but decreased to around 60 % of pre-starvation levels at 19 h after glucose removal (13).

Acute glucose depletion is also known to trigger a rapid drop in the cytosolic pH and subsequent disassembly of the vacuolar ATPase (V-ATPase), a proton pump required for the acidification of the vacuole. Shutdown of V-ATPase activity has been proposed to help preserve ATP by reducing energy consumption during starvation (43). It is possible that a similar principle applies to other ATP-dependent proteins, including chaperones and Hsp104 disaggregase (24), whereas specific ATP-dependent factors required for proteasomal degradation, such as the E1 ubiquitin activating enzyme and proteasomal ATPases appear to retain sufficient function under these conditions, as inferred from the sustained degradation of misfolded proteins observed in our study. Taken together, our data suggest that degradation-mediated PQC represents a prioritized cellular pathway under this form of nutrient stress.

ATP levels in mammalian cells are typically maintained within a narrow range under basal conditions; however they can fluctuate in response to metabolic stress, including hypoxia and nutrient limitation (44–46). Cell types with especially high energy demands, such as neurons, are particularly sensitive to disturbances in energy metabolism, and energy failure has been implicated in cellular dysfunction and neurodegenerative disease (47–48). Our findings that clearance of specific aggregation-prone proteins under metabolic stress increasingly relies on membrane contact site-associated pathways, likely because of limitations in factors other than the proteasome itself, may therefore provide insight into how proteostasis is maintained under acute energy limitation in mammalian systems. Although a direct mammalian counterpart of the yeast nucleus-vacuole junction has not been defined, mammalian cells contain ER-lysosome/ endolysosome membrane contact sites that may perform partly analogous functions (27).

In conclusion, our findings show that yeast cells preserve proteasome-mediated PQC during acute glucose starvation, with NVJ membrane contact sites supporting degradation of specific misfolded proteins during prolonged starvation. More broadly, these results demonstrate that yeast cells exhibit marked proteostasis resilience during acute glucose starvation, maintaining degradation-mediated PQC despite substantial fluctuations in cellular energy levels. Given the conservation of proteasome-mediated proteostasis and membrane contact site biology across eukaryotes, our findings raise the possibility that related mechanisms may also operate in mammalian cells to support protein quality control during acute metabolic stress.

## EXPERIMENTAL PROCEDURES

### Yeast strains

Yeast *Saccharomyces cerevisiae* strains used in this study are detailed in Table 1. All strains were isogenic to BY4741, except the *rpt2RF* strain, which was derived from SUB62. The description of strain construction is provided in Table 2. Strains were generated by homologous recombination, by transformation of recipient yeast strains with either plasmids linearized with the specified restriction enzymes, or with PCR-generated DNA constructs (Table 2), followed by the selection on the appropriate selective media. Molecular cloning followed standard protocols. The sequences of all primers used in this study are provided in Table 3. Successful genome integration of transformed constructs was confirmed by PCR.

**Table 1.**
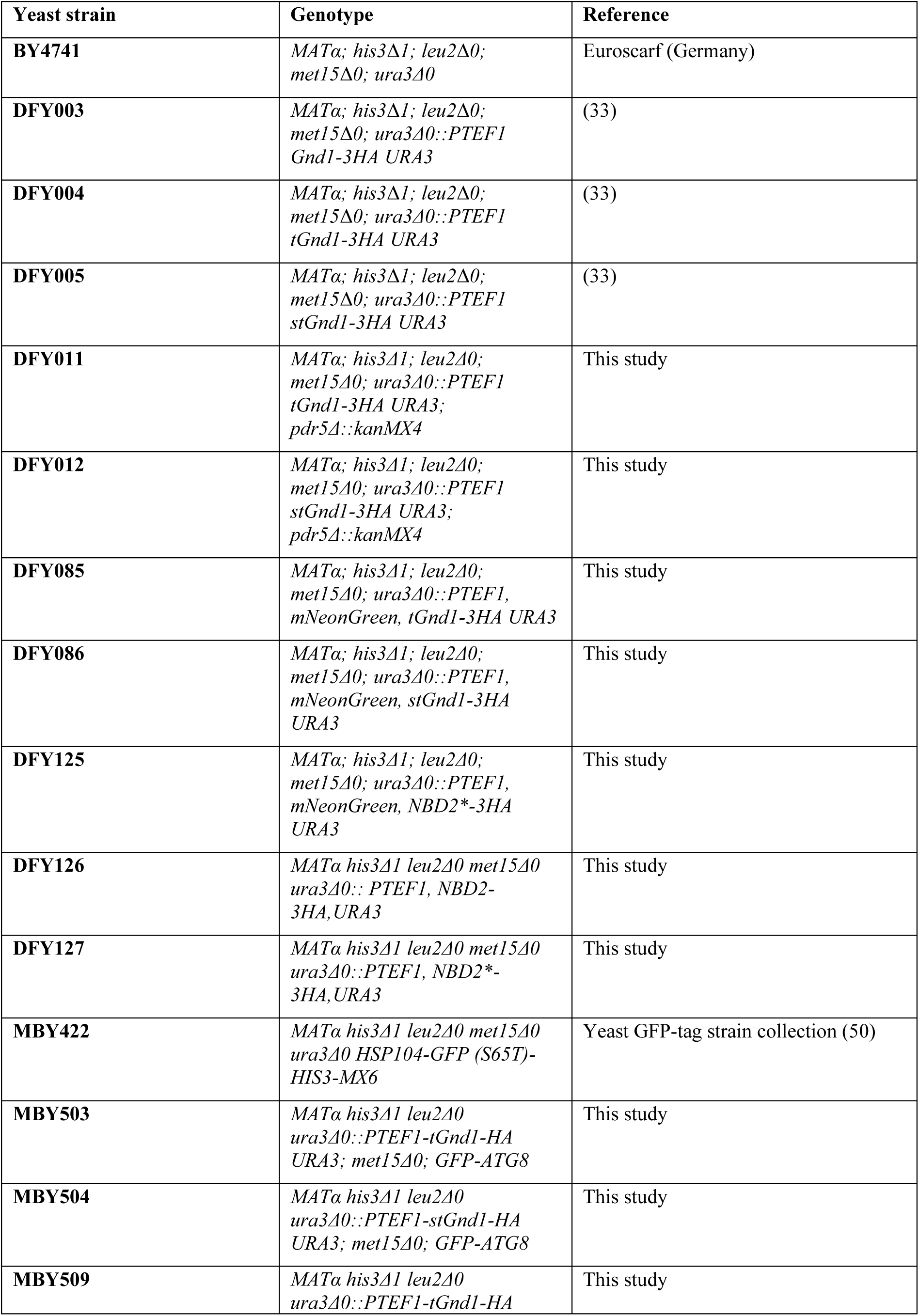
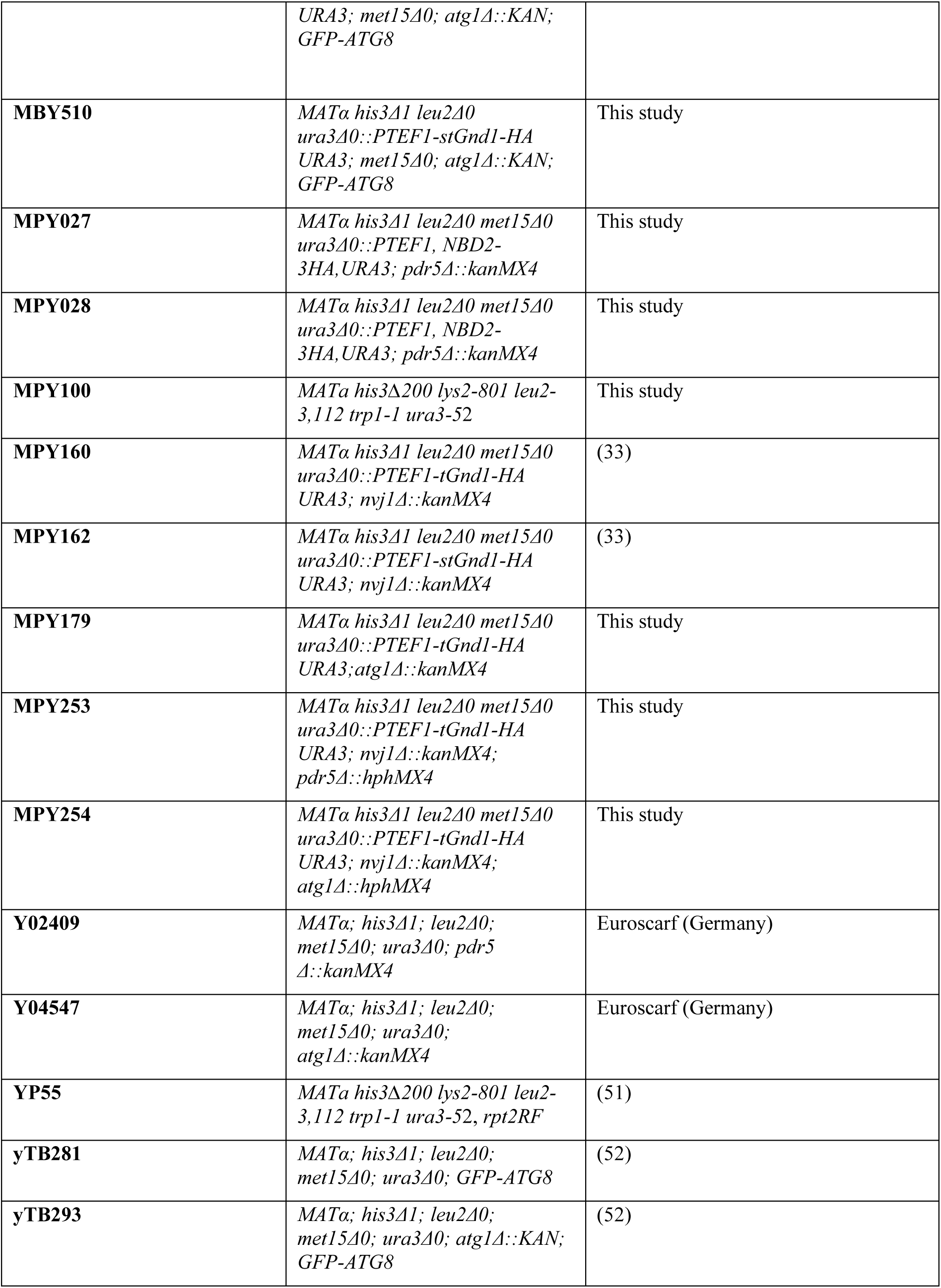
*Saccharomyces cerevisiae* strains used in this study.

**Table 2.** Yeast strain construction.

| Yeast strain constructed | Construction |  |
| --- | --- | --- |
|  | DNA used for transformation of the starting strain/restriction enzyme used to cut the DNA | Starting strain that was transformed with the indicated DNA |
| DFY011 | pDF040/AscI | Y02409 |
| DFY012 | pDF041/AscI | Y02409 |
| DFY085 | pDF081/AscI | BY4741 |
| DFY086 | pDF082/AscI | BY4741 |
| DFY125 | pDF106/AscI | BY4741 |
| DFY126 | pDF107/AscI | BY4741 |
| DFY127 | pDF108/AscI | BY4741 |
| MPY027 | pDF107/AscI | Y02409 |
| MPY028 | pDF108/AscI | Y02409 |
| MPY100 | *RPT2* gene was amplified by PCR using wild type BY4741 genomic DNA as template and primers prMB987 and 988. Following transformation into *rpt2RF* mutant yeast strain YP55, colonies carrying wild type *RPT2* gene were selected at 37 C. | YP55 |
| MPY179 | pDF040/AscI | Y04547 |
| MPY253 | *pdr5Δ::hphMX*<br>(PCR using pAG32 as template and primers prMP052 and 053) | MPY160 |
| MPY254 | *atg1Δ::hphMX*<br>(PCR using pAG32 as template, and primers prMP058 and 059) | MPY160 |
| MBY503 | pDF040/AscI | yTB281 |
| MBY504 | pDF041/AscI | yTB281 |
| MBY509 | pDF040/AscI | yTB293 |
| MBY510 | pDF041/AscI | yTB293 |

**Table 3.**
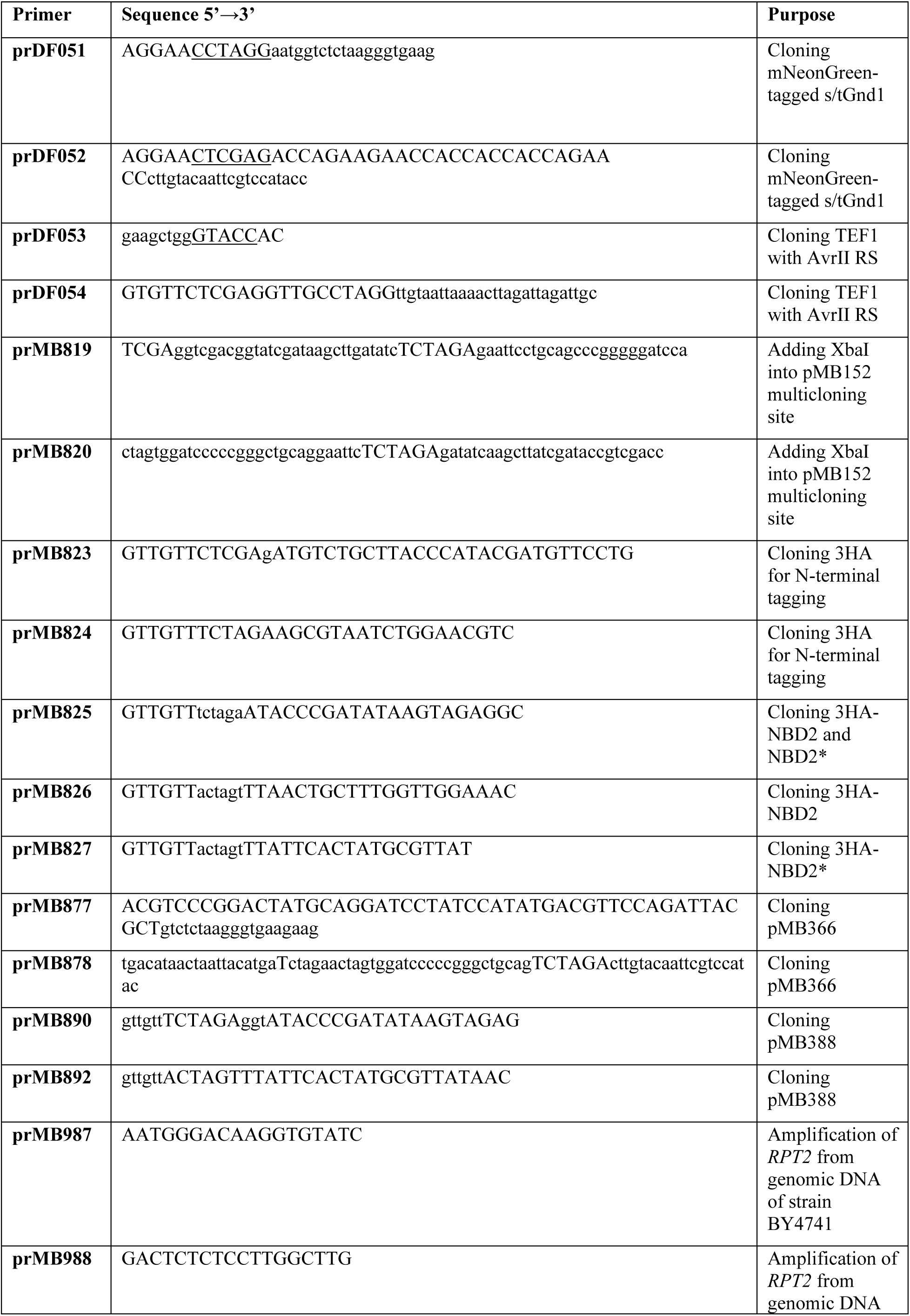
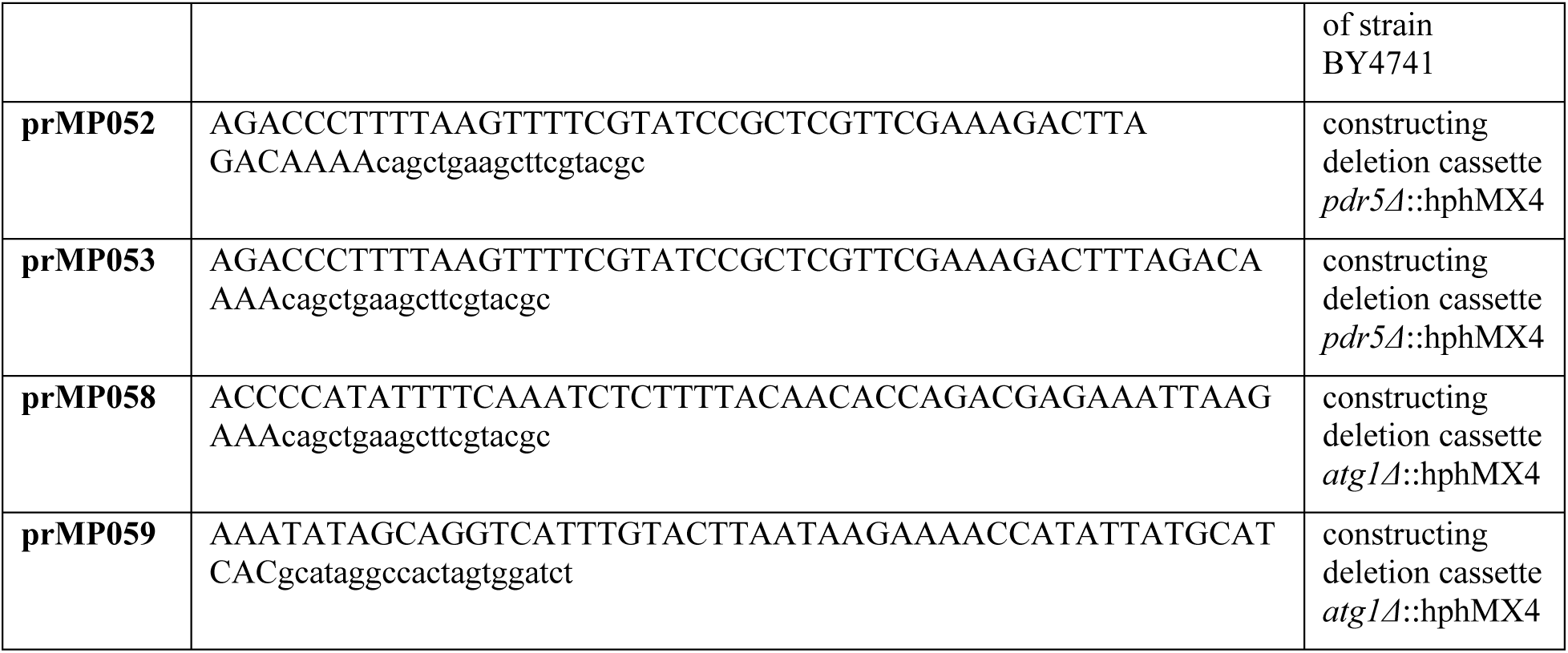
Primers used in this study.

### Plasmids

All plasmids used in this study are listed in Table 4. The description of the plasmid construction is listed in Table 5. Molecular cloning was performed by standard protocols. Sequences of primers used for construction are listed in Table 3.

**Table 4.**
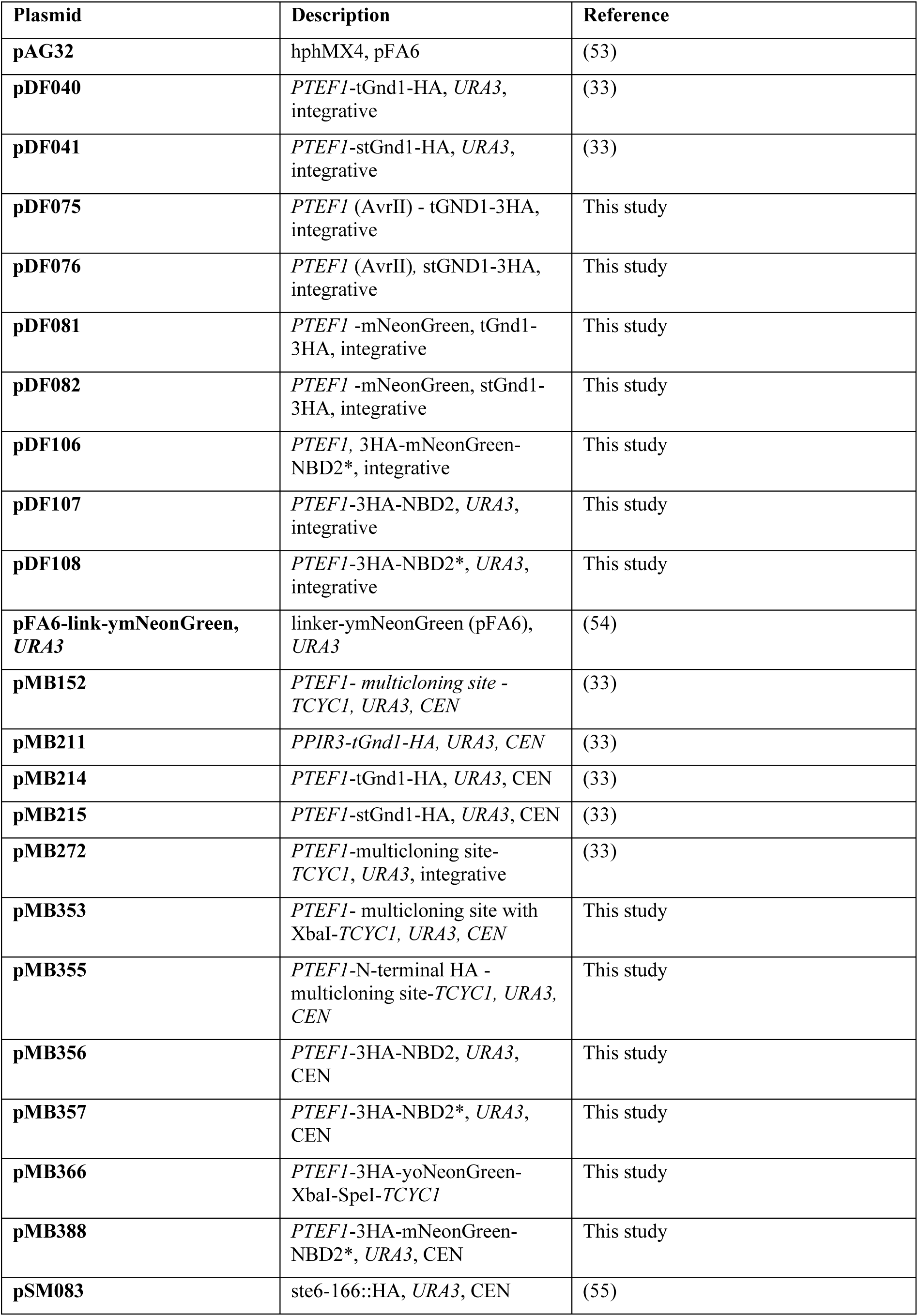
Plasmids used in this study.

**Table 5.** Plasmid construction.

| **Plasmid** | **Description of the construction** |
| --- | --- |
| pDF075,<br>pDF076 | To insert PTEF1 with AvrII restriction site at the N-terminus of stGnd1-HA and tGnd1-HA, the insert was amplified from pDF040 using primers prDF053 and prDF054, cut with XhoI/KpnI and ligated with similarly cut pDF040 and pDF041, yielding pDF075 or pDF076, respectively. |
| pDF081,<br>pDF082 | To insert mNeonGreen at the N-terminus of stGnd1-HA and tGnd1-HA, mNeonGreen was amplified from pFA6a-link-ymNeongreenSpHis5 using primers prDF051 and prDF052, cut with AvrII/XhoI and ligated with similarly cut pDF075 and pDF076, yielding pDF081 or pDF082, respectively. |
| pDF106,<br>pDF107,<br>pDF108 | To obtain integrative plasmids, 3HA-mNeonGreen-NBD2*, 3HA-NBD2, 3HA-NBD2* constructs from CEN plasmids pMB388, pMB356, pMB357 were cut with XhoI/SpeI and ligated with similarly cut pMB272, yielding pDF106, pDF107 or pDF108, respectively. |
| pMB353 | XhoI, SpeI-cut plasmid pMB152 was ligated with annealed primers prMB819 and prMB820. |
| pMB355 | XhoI, XbaI-cut plasmid pMB353 was ligated with similarly cut PCR amplified by primers prMB823 and prMB824 from plasmid pMB211 as a template. |
| pMB356 | SpeI, XbaI-cut plasmid pMB355 was ligated with similarly cut PCR amplified by primers prMB825 and prMB826 from wild type genomic DNA. |
| pMB357 | SpeI, XbaI-cut plasmid pMB355 was ligated with similarly cut PCR amplified by primers prMB825 and prMB827 from plasmid pSM083 as a template. |
| pMB366 | Plasmid was constructed by homologous recombination in yeast. PCR product amplified by primers prMB877, and prMB878 from plasmid pFA6-link-ymNeonGreen as a template was co-transformed to yeast with XbaI, EcoRI-cut plasmid pMB355, and selected for Ura<sup>+</sup> colonies. Plasmid was purified from yeast cells. |
| pMB388 | SpeI, XbaI-cut plasmid pMB366 was ligated with similarly cut PCR amplified by primers prMB890 and prMB892 using plasmid pMB357 as a template. |

### Yeast culture and growth media

Standard yeast culture media, such as yeast extract, peptone, dextrose (YPD) and ammonia-based synthetic complex dextrose (SCD) medium were used. YPD medium was prepared from 1% (w/v) yeast extract, 2% (w/v) peptone, and 2% (w/v) D-glucose (Formedium Ltd.). Media contained 2% D-glucose unless otherwise indicated. Antibiotic selections were made on solid YPD containing 300 mg/l hygromycin B. When grown in liquid media, culture tubes and baffled flasks with loose-fitting caps were used, and cells were incubated in an orbital shaker (Innova 40R, New Brunswick) with shaking at 240 rpm. Cells were grown at 30 °C unless indicated otherwise.

Acute glucose depletion was performed as follows. Overnight cultures grown in YPD were diluted in fresh YPD containing 2% glucose to an optical density OD_600_ of 0.2 and grown until reaching the mid-logarithmic phase (OD_600_ ≈ 0.6). Cells were then pelleted by centrifugation (3000 × g for 2 min at room temperature), washed once with sterile phosphate-buffered saline (PBS), and resuspended in YPD containing either 2%, 0.2% or 0.02% glucose to an optical density OD_600_ of 0.2. Cultures were incubated in the respective media for 90 min or 4 h, as indicated, prior downstream analysis. The analysis of yeast strains YP55 (*rpt2RF* mutant) and MPY100 (respective *RPT2* wild type), was performed as above, except that these strains were grown at 26°C overnight and to mid-logarithmic phase, and cultures were shifted to 30°C after resuspension of cells in fresh media containing different glucose concentrations.

### Measurement of ATP

Intracellular ATP was measured using the BacTiter-Glo assay (Promega) according to the manufacturer’s instructions. Briefly, 100 µL of yeast culture was mixed with an equal volume of BacTiter-Glo reagent, briefly agitated by using the integrated orbital shaking function of the GloMax® Explorer Multimode Microplate Reader (Promega, USA), followed by a 5-minute room-temperature incubation prior to luminescence measurement. Luminescent signal (relative luminescent units, RLU) was normalized to the optical density of yeast cultures (RLU/OD_600_).

### Cycloheximide chase and Western blot analysis

Cycloheximide (Sigma-Aldrich, St Louis, MO, USA) was added to the cell culture at a final concentration of 100 μg/ml to inhibit *de novo* protein synthesis. At the indicated time points after the addition of cycloheximide, 1.0 OD_600_ unit of cells was collected by centrifugation at 11,000 rpm for 4 min at 4°C, cell pellet was snap-frozen in liquid nitrogen, and stored at −20°C until samples from all time points had been collected.

Total cell lysates were prepared using an alkaline lysis protocol (49) with some modifications. Briefly, cell pellets were resuspended in 100 µL of dH_2_O, then 100 µL of ice-cold 0.2 M NaOH was added, followed by incubation on ice for 5 min. Samples were centrifuged at 11,000 rpm for 4 min at 4°C, pellets were resuspended in 50 µL sample buffer (0.06 M Tris-HCl pH 6.8, 5% glycerol, 2% SDS, 4% β-mercaptoethanol, and 0.0025% bromophenol blue), incubated at 97°C for 3 min, cooled briefly, and centrifuged again at 11,000 rpm for 5 min at 23°C. The resulting supernatants containing total cell lysates were collected.

Total cell lysates were resolved by SDS-PAGE and transferred to Immobilon®-FL PVDF Membrane (Millipore, Bedford, MA, USA) for immunoblotting. Primary antibodies used were mouse monoclonal anti-HA (clone 12CA5, 1:1000; Ogris Laboratory, Max F. Perutz Laboratories, Vienna, Austria), and rabbit polyclonal anti-GFP (G1544, 1:1000; Sigma Aldrich, USA). Signals were detected using HRP-conjugated anti-mouse (#7076, 1:2000) or anti-rabbit (#7074, 1:1000) secondary antibodies (Cell Signaling Technology). Chemiluminescent signals were captured with a ChemiDoc MP Imaging System and quantified via ImageLab software (Bio-Rad Laboratories). Total proteins were visualized using stain-free imaging technology (BioRad) and used as a loading control. For quantitative analysis of protein levels, intensity of the band corresponding to the protein of interest was normalized to the intensity of the total protein signal from the respective lane. Data are presented as the mean ± S.D. of at least two independent experiments.

### Fluorescence microscopy

Cells were fixed in 0.8% (v/v) formaldehyde for 10 min at room temperature, harvested by centrifugation (8000 RPM, 1 min, RT), and washed twice with phosphate-buffered saline (PBS). The resulting cell pellets were resuspended in PBS and applied to coverslips pre-coated with 1 mg/mL concanavalin A (Sigma-Aldrich) to promote attachment of cells. Images were acquired using an Olympus FV3000 laser-scanning confocal microscope (Olympus) equipped with an Olympus-DP74 digital camera. Samples were visualized using a 60× oil-immersion objective (Olympus UPlanSApo 60x/1.35 Oil Microscope Objective). Image acquisition was controlled by the FV31S-SW Fluoview software. Images were captured as complete Z-stacks spanning the entire volume of the cells. Final image analysis and figure preparation were performed using Fiji/ImageJ.

### Statistical analysis

All data are presented as the mean ± S.D. derived from the indicated number of independent biological replicates (n ≥ 2).

## DATA AVAILABILITY

All data are contained within the manuscript.

## SUPPORTING INFORMATION

This article contains supporting information (Figure S1, Figure S2, Figure S3).

## ACKNOWLEDGMENTS

Strains yTB281 and yTB293 were a kind gift from Claudine Kraft. Strain YP55 was a kind gift from Elah Pick. Plasmid pFA6-link-ymNeonGreen was a gift from Bas Teusink via Addgene.

## ABBREVIATIONS

The abbreviations used are: NVJ, nucleus-vacuole junctions; PQC, protein quality control.

## Supporting information

– **Supplementary figures S1 – S3**

**Supplementary figure S1.**
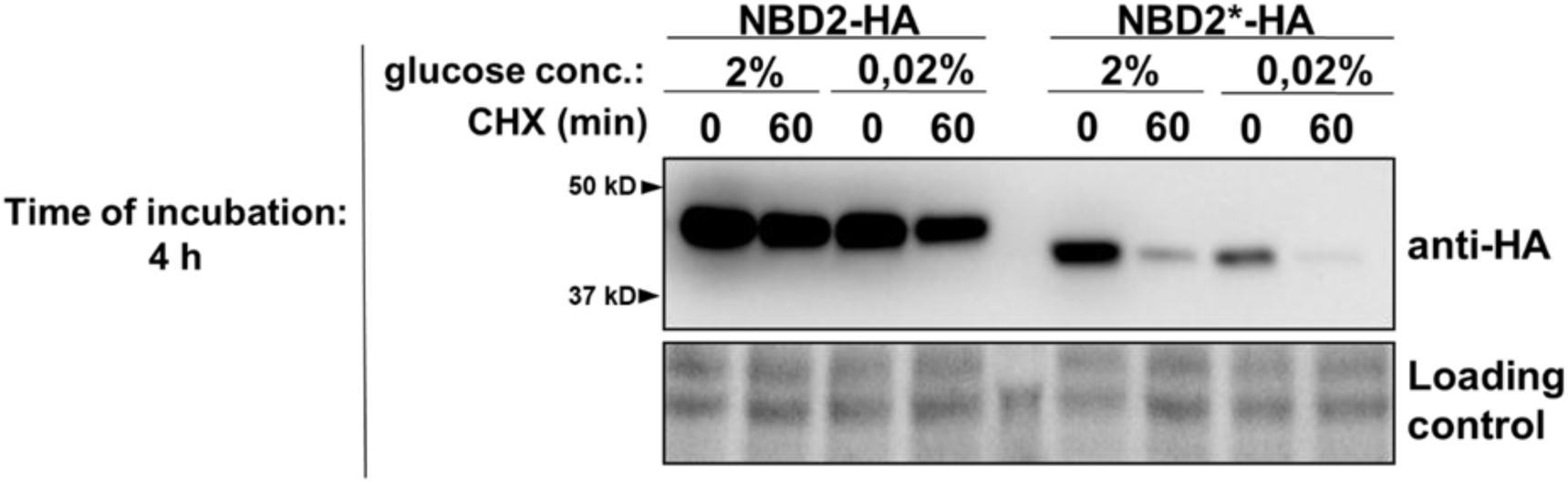
Selective degradation of the misfolded protein NBD2*-HA is maintained during acute glucose starvation. Representative immunoblots of native NBD2-HA and the misfolded variant NBD2*-HA. Exponentially growing yeast cells expressing NBD2-HA or NBD2*-HA were subjected to acute glucose depletion (0.02% glucose) or glucose replete conditions (2% glucose) as a control, for 4 h. Protein stability was assessed by cycloheximide chase assay (CHX, 100 µg/mL) by analyzing protein levels at indicated time points by Western blot (anti-HA). Stain-free total protein is shown as a loading control.

**Supplementary figure S2.**
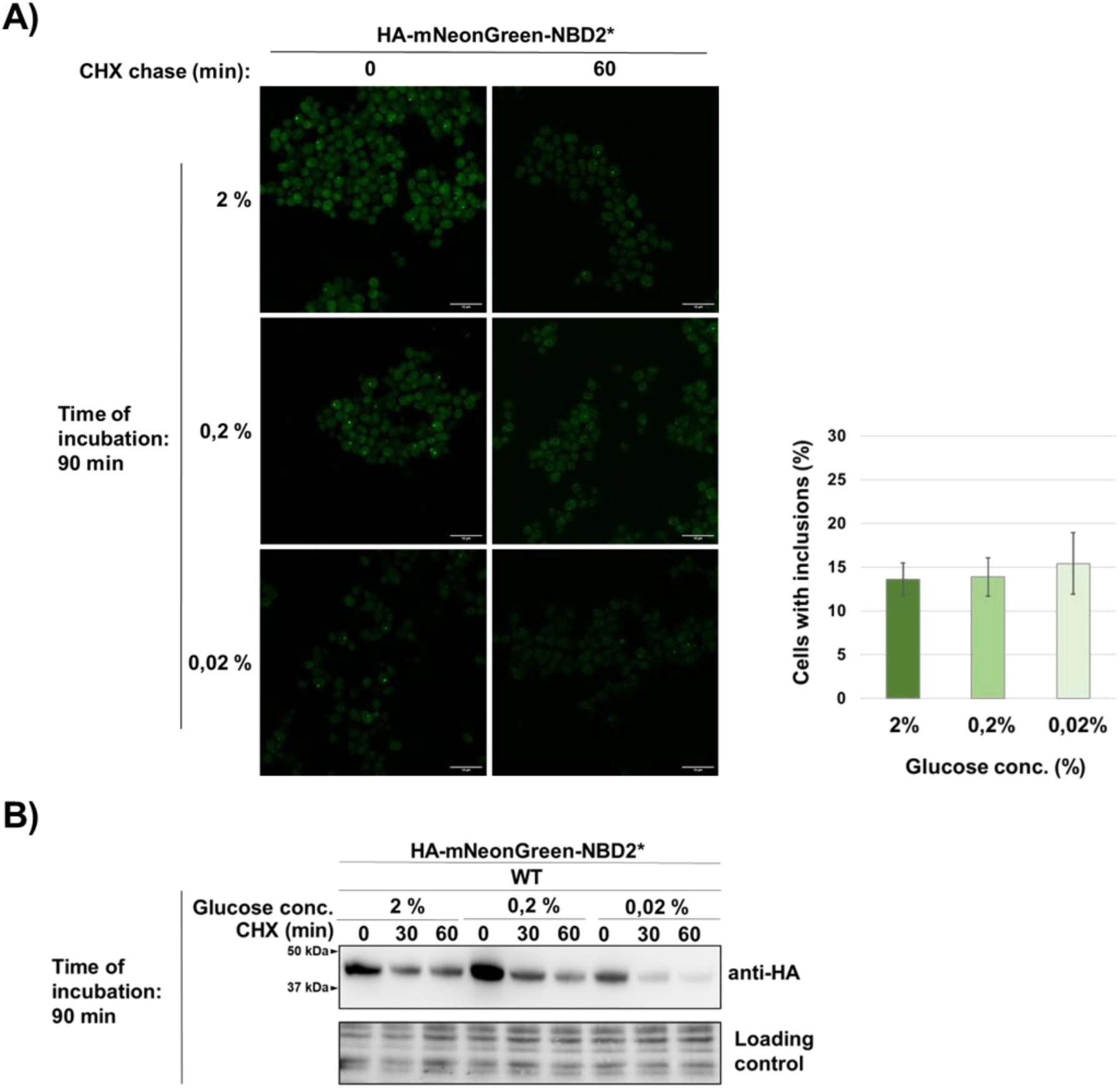
Selective degradation of HA-mNeonGreen-NBD2* persists during acute glucose starvation in *S. cerevisiae*. **A**, Representative images of yeast cells expressing HA-mNeonGreen-NBD2* under glucose-replete conditions (media containing 2% glucose) or acute glucose starvation (media containing 0.02% glucose) for 90 min. Yeast cells expressing HA-mNeonGreen-NBD2* were exponentially grown in the medium containing 2% glucose, and either maintained under glucose replete conditions (2% glucose) or subjected to acute glucose depletion (0.02% glucose) for 90 min prior to imaging by confocal microscopy. Images show whole Z-stacks. Scale bar, 10 µm. The graph shows the percentage of cells that contain inclusions. Data are expressed as mean ± SD from two independent experiments. **B**, Representative immunoblots showing turnover of the model misfolded protein HA-mNeonGreen-NBD2*. Cells grown as in (A) were subjected to cycloheximide chase assay (CHX, 100 µg/mL), harvested at indicated time points after cycloheximide addition and protein levels were analyzed by Western blot (anti-HA). Stain-free total protein is shown as a loading control.

**Supplementary figure S3.**
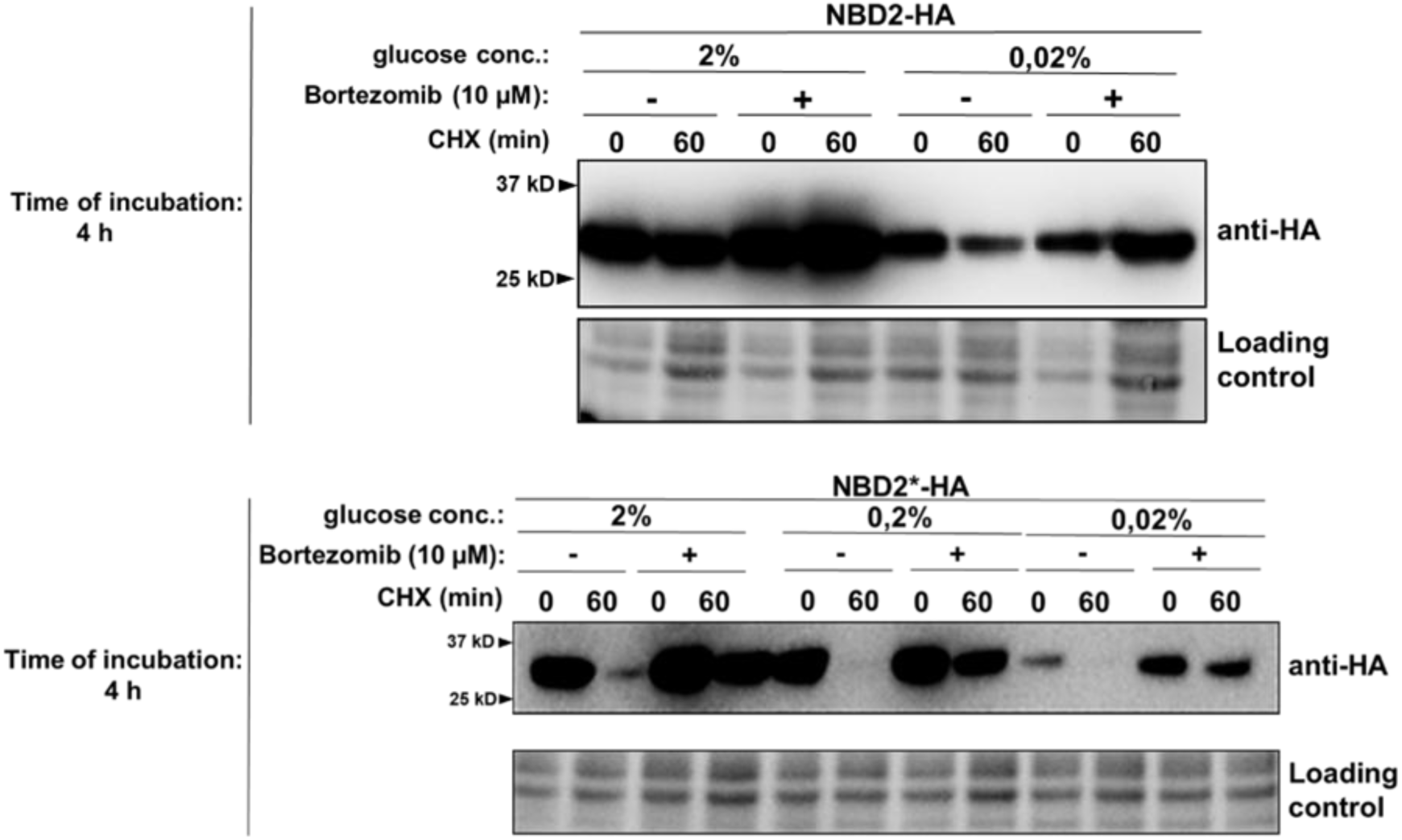
Degradation of NBD2*-HA during acute glucose starvation depends on proteasomal proteolytic activity. Representative immunoblots of NBD2-HA and NBD2*-HA in the presence or absence of proteasomal inhibition. Exponentially growing yeast cells expressing NBD2-HA or NBD2*-HA were subjected to acute glucose depletion (0.02% glucose) or glucose replete conditions (2% glucose) as a control, for 4 h. Protein stability was assessed by cycloheximide chase assay (CHX, 100 µg/mL) by analyzing protein levels at indicated time points by Western blot (anti-HA). Proteasome dependency was evaluated by treating cells with proteasomal inhibitor bortezomib (10 µM) for 30 min before cycloheximide addition. Stain-free total protein is shown as a loading control.

